# RNA-mediated Allosteric Activation in CRISPR-Cas13a

**DOI:** 10.1101/2023.07.27.550797

**Authors:** Souvik Sinha, Adrian M. Molina Vargas, Pablo R. Arantes, Amun Patel, Mitchell R. O’Connell, Giulia Palermo

## Abstract

Cas13a is a recent addition to the CRISPR-Cas toolkit that exclusively targets RNA, which makes it a promising tool for RNA detection. The protein uses a CRISPR RNA (crRNA) to target RNA sequences, which are cleaved by a composite active site formed by two ‘Higher Eukaryotes and Prokaryotes Nucleotide’ (HEPN) catalytic domains. In this system, an intriguing form of allosteric communication controls RNA cleavage activity, yet its molecular details are unknown. Here, multiple-microsecond molecular dynamics simulations are combined with graph theory and RNA cleavage assays to decipher this activation mechanism. We show that the binding of a target RNA acts as an allosteric effector of the spatially distant HEPN catalytic cleft, by amplifying the allosteric signals over the dynamical noise, that passes through the buried HEPN interface. Critical residues in this region – N378, R973, and R377 – rearrange their interactions upon target RNA binding, and alter allosteric signalling. Alanine mutation of these residues is experimentally shown to select target RNA over an extended complementary sequence beyond guide-target duplex, for RNA cleavage. Altogether, our findings offer a fundamental understanding of the Cas13a mechanism of action and pave new avenues for the development of more selective RNA-based cleavage and detection tools.

## INTRODUCTION

CRISPR (Clustered Regularly Interspaced Short Palindromic Repeats) and their associated (Cas) proteins are RNA-guided prokaryotic adaptive immune systems that protect bacteria against invading genetic elements (1). The transformative use of CRISPR-Cas9 for genome editing (2) led to the “boom” of the “CRISPR field” and the discovery of novel CRISPR-Cas systems that remarkably expand applications in genome editing and beyond (3–5). Cas13a (formerly known as C2c2) is a recently discovered CRISPR-associated protein that targets RNA (6, 7), a property that is powerful for RNA detection, regulation, and imaging (8–13). The Cas13a effector protein complexes with a CRISPR RNA (crRNA) sequence, which is used as a guide to form the RNA-targeting interference complex (6, 7, 14). The latter can cleave single-stranded RNA (ssRNA) sequences using two Higher Eukaryotes and Prokaryotes Nucleotide (HEPN) catalytic domains (7, 15), which are found in RNA targeting enzymes (16). Upon activation of the HEPN domains, Cas13a degrades its target RNAs through *cis* cleavage and other solvent-exposed ssRNAs through a non-specific *trans* cleavage activity (6, 7). This is a potent “collateral damage” that has enabled the development of ultrasensitive RNA detection tools like SHERLOCK (13), SPRINT (12), direct SARS-CoV-2 RNA detection assays (17), and many others.

The biophysical function of the Cas13a effector is characterized by intricate allosteric signalling, whose molecular details are highly unclear (18). Structures of Cas13a from *Leptotrichia buccalis* (Lbu) reveal a bilobed architecture, comprising a “REC” ⍺-helical lobe that recognizes the crRNA and a “NUC” lobe holding the catalytic HEPN1-2 domains and a helical Linker (**Figure 1**) (19). Biochemical studies have shown that the binding of a complementary target RNA (tgRNA) to the REC lobe allosterically activates the HEPN domains (18), which are spatially distant, to form a composite active site for RNA cleavage (7, 15). However, the mechanism of this activation and information transfer from the tgRNA binding site to the HEPN1-2 catalytic cleft is poorly understood.

**Figure 1.**
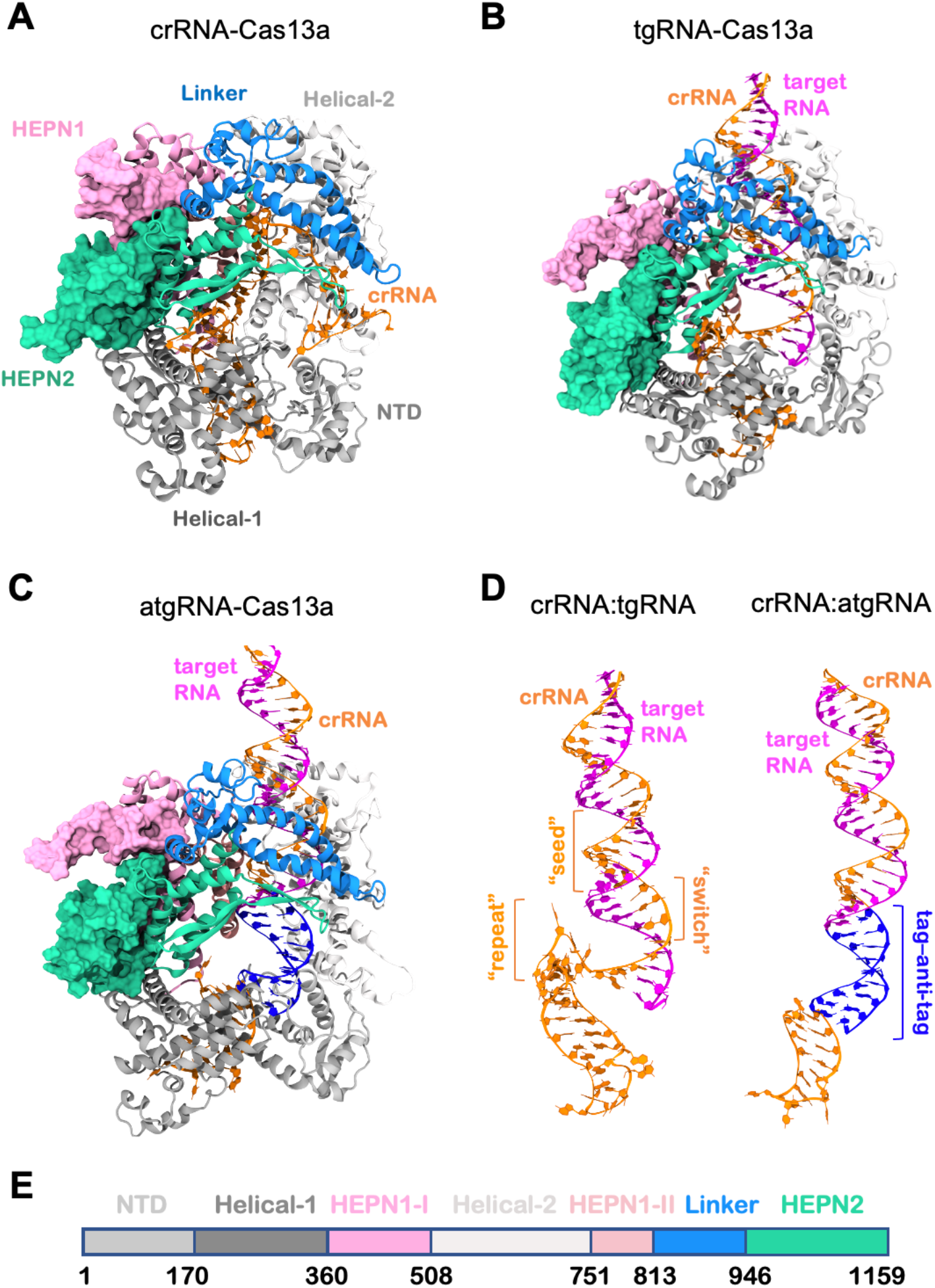
Overview of the RNA-bound Cas13a complexes from *Leptotrichia buccalis*. **A-C.** Structures of the *Leptotrichia buccalis* Cas13a bound to a guide CRISPR RNA (crRNA, A), a target RNA (tgRNA, B), and a tag–anti-tag RNA (atgRNA, C) (19). The Cas13a protein is shown as cartoons, using a molecular surface representation for the HEPN1 (mauve) and HEPN2 (green) catalytic domains. **D.** Structure of the RNA in the tgRNA-(left) and atgRNA-(right) bound systems. The crRNA (orange) forms a duplex with the tgRNA (magenta), whose complementarity is extended at the 5’ flank of the crRNA through a tag–anti-tag pairing (blue). The “seed” (nucleotides, nt. 9-14) and “switch” (nt. 5-6) regions of the crRNA, as well as the crRNA repeat region (nt. (A(−5)–(C-1)), are also indicated. **E.** Protein sequence and domain colour code.

Interestingly, the crRNA spacer – i.e., the 28 nucleotides (nt.) sequence that guides binding of the tgRNA – was shown to hold a critical role in the onset of the allosteric response (18). The spacer contains a “seed” region (nt. 9-14), where perfect base pairing is required for tgRNA binding, and a “switch” region (nt. 5-8, **Figure 1D**), whose binding of the tgRNA has been suggested to induce the activation of the catalytic cleft (18). Nevertheless, the allosteric response of the “seed” and “switch” regions toward activation is unclear.

Functional studies, both *in-vitro* and *in-vivo*, have also shown that extending the duplex complementarity at the 5’ flank of the crRNA spacer, impacts the RNA degrading ability of Cas13a (20). Such extended pairing beyond the guide-target duplex is called tag (segment from the crRNA) – anti-tag (segment from the tgRNA) pairing (blue segment in **Figure 1D**). This dependence on the complementarity length was proposed as a strategy to discriminate between self- and non-self-targets, which is key for Cas13-based technologies (21). This suggests the allosteric response is regulated by the degree of complementarity between the guide crRNA (also referred to as a guide-RNA) and tgRNA.

In light of this knowledge, understanding the allosteric regulation of Cas13a activation and the role of RNA is crucial to improve the RNA targeting specificity of Cas13a, and engineering its controlled function to discriminate non-self-targets.

Here, we used extensive molecular dynamics (MD) simulations, reaching a conformational sampling of ∼170 μs, and conducted fluorescent RNA trans-cleavage assays, providing a comprehensive RNA-mediated allosteric mechanism of the Cas13a protein. Most significantly, our computational investigations disclose critical residues whose mutation in alanine is shown experimentally to select tgRNA over an extended tag–anti-tag complementarity, for RNA cleavages. These findings offer a fundamental understanding of the Cas13a mechanism of action and pave new avenues for the development of more selective RNA-targeting CRISPR-Cas13a systems.

## MATERIAL AND METHODS

### Structural models

Molecular simulations were based on the structure of the *Leptotrichia buccalis* (Lbu) Cas13a bound to a crRNA (PDB: 5XWY, at 3.2 Å resolution (15)), obtained via cryo-EM, and on the structure of the LbuCas13a in complex with a crRNA and tgRNA (PDB: 5XWP), obtained by single-wavelength anomalous diffraction at 3.08 Å resolution (15). The LbuCas13a bound to an extended tag–anti-tag RNA (atgRNA) was built by including a longer duplex with eight base-pair extended atgRNA, obtained from the cryo-EM structure of the *Leptotrichia shahii* Cas13a bound to atgRNA (PDB: 7DMQ, at 3.06 Å resolution) (20). In all systems, we reinstated the catalytic H1053 and R1048 in the HEPN domains, which were mutated in alanine in the experimental structures (15). The systems were solvated, leading to simulation cells of ∼ 138 * 97 * 130 Å^3^, and neutralized by the addition of an adequate number of Na^+^ ions.

### Molecular dynamics simulations

Molecular dynamics (MD) simulations were performed using a simulation protocol tailored for protein/nucleic acid complexes. We employed the Amber ff19SB (22) force field, including the χOL3 corrections for RNA (23, 24). The TIP3P model was used for explicit water molecules (25). Production runs were carried out in the NVT ensemble, using an integration time step of 2 fs. For each system, i.e., Cas13a bound to a crRNA (crRNA-Cas13a), in complex with a crRNA and tgRNA (tgRNA-Cas13a) and bound to an extended tag–anti-tag RNA (atgRNA-Cas13a), we performed ∼5 μs of MD simulations in three replicates. We also carried out a ∼5 μs long simulation of a tgRNA-bound complex substituting the A(−3) base of the crRNA with cytosine C(−3). Subsequently, we considered four variants (R377A, N378A, R963A, and R973A) of Cas13a bound to a tgRNA, and including an atgRNA, similar to the wild-type (WT) Cas13a complexes. These systems were also simulated for ∼5 μs in three replicates. Overall, we accumulated ∼165 μs of total sampling. All simulations were performed using the GPU-empowered version of the AMBER 20 code (26). Analyses were performed over the aggregated multi-μs sampling collected for each complex (i.e., ∼15 μs). This was motivated by our previous studies of allostery in CRISPR-Cas systems (27–31), showing that an aggregated multi-μs sampling offers a robust ensemble for the purposes of our analysis (described below).

### Analysis of Jensen-Shannon Distances

To characterize the difference in the conformational dynamics of the HEPN domains, we analysed the distributions of all intra-backbone dihedrals (BB torsions) and backbone C_α_ distances (BB distances) in the investigated systems. To compare the distributions of the abovementioned features between any two of our systems, we computed the Jensen-Shannon Distance (*JSD* or *D*_*JS*_), a symmetrized version of Kullback-Leibler divergence (*D*_*KL*_).(32) The *JSD* ranges from 0 to 1, where 0 corresponds to two identical distributions and 1 corresponds to a pair of separated distributions. For two distributions *P*_*i*_and *P*_*j*_, and considering a feature *x*_*f*_ from two different ensembles *i* and *j*,

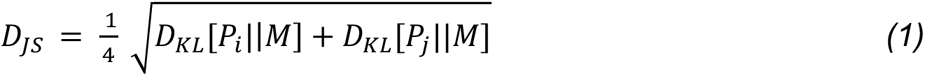

where 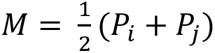. The Kullback-Leibler divergence, *D*_*KL*_, corresponds to two distributions *P*_*i*_ and *P*_*j*_ is of the following form:

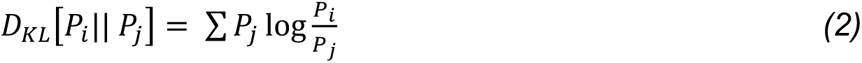

*JSD* values were computed using the Python ENSemble Analysis (PENSA) open-source library (33). Kernel density estimations of the *JSD* values were plotted to describe the *JSD* range and compare the systems.

### Inter-domain correlations

To evaluate the inter-dependent coupling between the Cas13a domains and the nucleic acids, we computed inter-domain generalized correlation scores (34). For each protein residue *i*, a correlation score (*Cs*) parameter can be computed as:

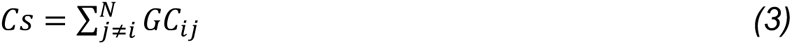

which sums the atom-based generalized correlations, *GC*_*ij*_, established by residue *i* with the residues *j*, based on Shannon’s entropy estimation of mutual information (35). *Cs* are a measure of the number and the intensity of the *GC*_*ij*_coefficients displayed by each residue pair. To detail inter-domain correlations, the *Cs* were accumulated and normalized. First, the *Cs* were calculated for each residue *i* belonging to a specific protein domain (e.g., HEPN1(I)), with the residues *j* belonging to another protein domain of interest (e.g., HEPN2). Then, the *Cs* were accumulated over all residues *j* of each specific domain and normalized by the number of coupling residues. This resulted in a set of inter-domain *Cs*, ranging from 0 (not-correlated) to 1 (correlated), measuring the strength of the overall correlation that each domain establishes with the others.

### Dynamic network and Signal-to-Noise Ratio

To characterize the allosteric pathways of communication, network analysis was applied (36). In dynamical networks, C⍺ atoms of proteins and backbone P atoms of nucleotides, as well as N1 atoms in purines, and N9 in pyrimidines, are represented as nodes, connected by edges weighted by the generalized correlations *GC*_*ij*_ according to:

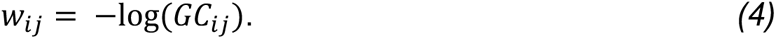

From the dynamical network, we estimated the efficiency of crosstalk between the crRNA spacer regions (i.e., “seed” (nt. 9-14); “switch” (nt. 5-8); as well as nt. 1-4; nt. 15-18 and nt. 19-22) and the catalytic residues (R472, H477, R1048, H1053) through a Signal-to-Noise Ratio (*SNR*) measure. *SNR* measures the preference of communication between predefined distant sites – i.e., the signal – over the remaining pathways in the network – i.e., the noise, estimating how allosteric pathways stand out (i.e., are favourable) over the entire communication network.

For the *SNR* calculation, we first computed the optimal (i.e., the shortest) and top five sub-optimal pathways (with longer lengths, ranked compared to the optimal path length) between all crRNA bases and the Cas13a residues, using well-established algorithms (*vide infra*). Then, the cumulative betweennesses of each pathway (*S*_*k*_) was calculated as the sum of the betweennesses of all the edges in that specific pathway:

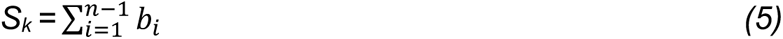

where *b*_*i*_ is the edge betweenness (i.e., the number of shortest pathways that cross the edge, measuring the “traffic” passing through them) between node *i* and *i* + 1, and *n* is the number of edges in the *K*^*th*^ pathway. The distribution of *S*_*k*_ between the crRNA bases and all protein residues was defined as the noise, whereas the distribution of *S*_*k*_ between the crRNA nucleotide regions of interest (e.g., “seed”) and the HEPN1-2 catalytic residues were considered as signals. Since *S*_*k*_ depends on the number of edges present in the path, to ensure consistency, we characterized the *SNR* on the basis of shorter (edge count: 6-8), medium (edge count: 9-11), and longer (edge count: 12-14) paths. Notably, while the optimal path corresponds to the most likely mode of communication, suboptimal paths can also be crucial routes for communication transfer (29, 36). Hence, in addition to the optimal path, we also considered the top five sub-optimal pathways for our *SNR* analysis. Well-established algorithms were employed for shortest-path analysis. The Floyd-Warshall algorithm was utilized to compute the optimal paths between the network nodes. The five sub-optimal paths were computed in rank from the shortest to the longest, using Yen’s algorithm, which computes single-source *K*-shortest loop-less paths (i.e., without repeated nodes) for a graph with non-negative edge weights (37). To identify residues important for allosteric communication, we computed the occurrence of each residue appearing in at least one of the pathways (i.e., optimal and sub-optimal). This analysis also reports on the conservation of allosteric pathways, as pathways characterized by a lesser number of residues with higher occurrence are likely to be more conserved than those exhibiting a greater number of residues with lower occurrence.

Finally, the *SNR* corresponding to signals from each crRNA region (i.e., “seed” (nt. 9-14); “switch” (nt. 5-8); as well as nt 1-4; nt. 15-18 and nt. 19-22) to the HEPN1-2 catalytic core residues (R472, H477, R1048, H1053) was computed as:

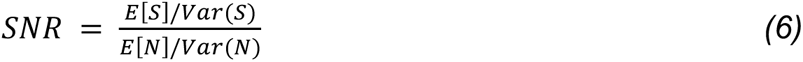

where *E*(*S*) and *Var*(*S*) correspond to the expectation and variance of the signal distribution respectively; and *E*(*N*) / *Var*(*N*) are the expectation/variance of the noise distribution. To provide the significance of the signal over the noise, we used a general approach based on *p*-value calculation. Our goal was to test the hypothesis that the signal is an outlier of the noise distribution. We can construct a best-fit probability distribution based on the noise data by treating our variable (the sum of betweennesses) as stochastic. Treating the signal as a sample from this population, we can assess the rarity of that sample’s mean when randomly collecting samples of the same size. This rarity is defined as the *p*-value of the sample. The *p*-value is then computed using the *Z*-score:

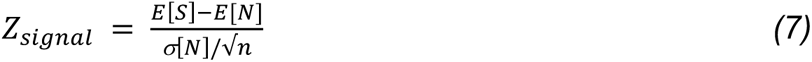

where *σ*[*N*] corresponds to the standard deviation of the noise, and *n* is the number of signals. Here, we found that the best-fit distributions for the noise follow a log-normal distribution. Hence, we took the logarithm of the signals and noise to transform the data points into a normal distribution. From this transformed distribution, we obtained the mean and standard deviations for the calculation of the *Z*_*signal*_. All networks were built using the Dynetan Python library (36). Path-based analyses were performed using NetworkX Python library (38) and our in-house Python scripts.

### Cas13a protein expression and purification

For expression of wild-type LbuCas13a we used Addgene Plasmid #83482 (7). LbuCas13a mutants were cloned from the WT vector via site-directed mutagenesis using the primers indicated in Table S1.

All constructs were purified as previously described (7, 40), with some modifications. Briefly, expression vectors were transformed into Rosetta2 DE3 grown in LB media supplemented with 0.5*i* w/v glucose at 37 °C. Protein expression was induced at mid-log phase (OD_600_ ∼0.6) with 0.5 mM IPTG, followed by incubation at 16 °C overnight. Cell pellets were resuspended in lysis buffer (50 mM HEPES [pH 7.0], 1 M NaCl, 5 mM imidazole, 5*i* (v/v) glycerol, 1 mM DTT, 0.5 mM PMSF, EDTA-free protease inhibitor [Roche]), lysed by sonication, and clarified by centrifugation at 15,000g. Soluble His_6_-MBP-TEV-Cas13a was isolated over metal ion affinity chromatography, and in order to cleave off the His_6_-MBP tag, the protein-containing eluate was incubated with TEV protease at 4 °C overnight while dialyzing into ion exchange buffer (50 mM HEPES [pH 7.0], 250 mM NaCl, 5*i* (v/v) glycerol, 1 mM DTT). The cleaved protein was loaded onto a HiTrap SP column (GE Healthcare) and eluted over a linear KCl (0.25–1 M) gradient. Fractions containing LbuCas13a were pooled, concentrated, and further purified via size-exclusion chromatography on an S200 column (GE Healthcare) in gel filtration buffer (20 mM HEPES [pH 7.0], 200 mM KCl, 5*i* glycerol (v/v), 1 mM DTT), snap-frozen in liquid N_2_ and were subsequently stored at −80°C. Protein purity was assessed by loading ∼2 ug of total protein in a 4-12*i* Bis-Tris gel and staining with Coomassie Blue dye.

### Fluorescent ssRNA nuclease assays

Mature crRNAs and ssRNA targets were commercially synthesized (IDT; Table S2). Cas13 trans-cleavage nuclease activity assays were performed in 10 mM HEPES pH 7.0, 50 mM KCl, 5 mM MgCl2, and 5*i* glycerol. Briefly, 100 nM LbuCas13a:crRNA complexes were assembled in for 30 min at 37°C. 100 nM of Rnase Alert reporter (IDT) and various final concentrations of ssRNA-target were added to initiate the reaction. These reactions were incubated in a fluorescence plate reader (Tecan Spark) for up to 60 min at 37°C with fluorescence measurements taken every 5 min (λ_ex_: 485 nm; λ_em_: 535 nm). Time-course and end-point values at 1 hour were background-subtracted, normalized, and analysed with their associated standard errors using GraphPad Prism9.

## RESULTS

### Conformational changes of the HEPN1-2 domains

MD simulations were carried out on three RNA-bound complexes of the LbuCas13a: the binary complex bound to a guide crRNA only (crRNA-Cas13a), and the ternary complexes in which Cas13a binds to a 28 nt. tgRNA matching the crRNA (tgRNA-Cas13a), and a 36 nt. Tag–anti-tag RNA (atgRNA-Cas13a, **Figure 1**). In these systems, we reinstated the catalytic H1053 and R1048 in the HEPN domains, which were mutated to alanine in the structures to prevent RNA cleavage (19). An aggregate sampling of ∼15 μs was collected for each system, from three simulation replicas of ∼5 μs each.

To characterize the difference in the HEPN1-2 conformational dynamics among the systems studied, we first analysed the distribution of the Jensen-Shannon Distances (JSD) (32), measuring the similarity of two distributions, ranging from 0 (similar) to 1 (dissimilar). The JSD distributions were computed to compare all intra-backbone distances (BB distances) and backbone torsions (BB torsions) of the HEPN domains in the investigated systems (see *Material and Methods*). We observe that, while overall the JSD distributions are similar for the overall HEPN1-2 domains, the backbone dynamics of the catalytic cleft (residues 470-480, 1045-1055) clearly display a separation between the crRNA- and tgRNA-Cas13a systems (**Figure 2A-B**, red distribution). A separation in the dynamics of the catalytic cleft is also observed when comparing the crRNA- and atgRNA-bound Cas13a (blue distribution), while the catalytic cleft dynamics in the atgRNA-Cas13a are least separated from those of the tgRNA-Cas13a (grey distribution). This observation is complemented by the Solvent Accessible Surface Area (SASA), showing that the HEPN1-2 catalytic cleft is significantly more accessible to the solvent in the tgRNA-bound system than in the crRNA-Cas13a binary system (**Figure 2C**), also deviating from the experimental structures (19). This indicates a closed catalytic core in the crRNA-bound complex, which opens in the tgRNA-Cas13a complex (**Figure 2D**). Interestingly, in the presence of an atgRNA, the catalytic cleft samples both open and closed conformations, also accessing states where the pocket is less accessible than in the crRNA-Cas13a.

**Figure 2.**
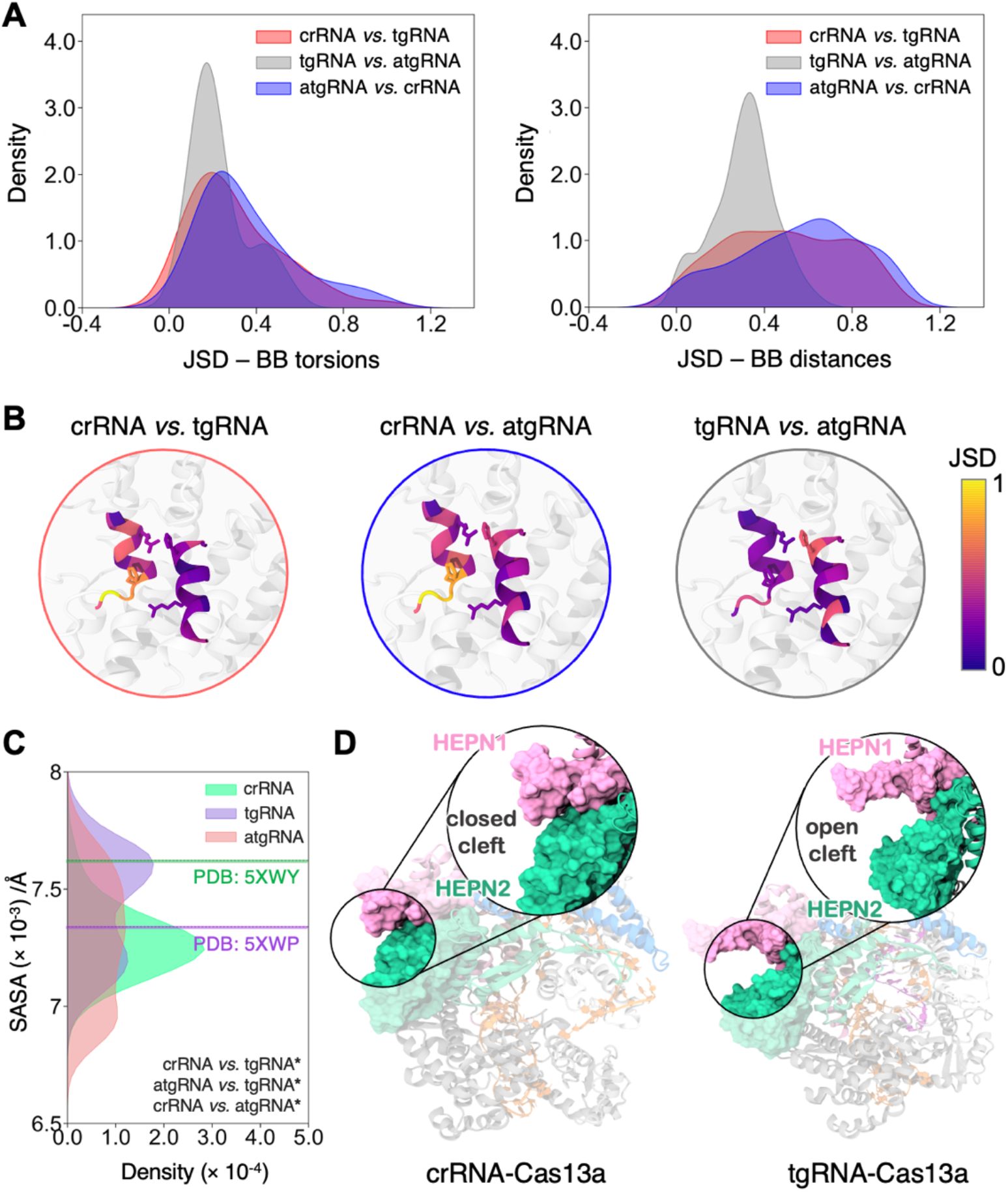
Structural dynamics of the HEPN1 and HEPN2 catalytic domains. **A.** Distribution of Jensen-Shannon Distances (JSD), comparing intra-backbone distances (BB distances) and backbone torsions (BB torsions) of the HEPN1-2 catalytic core helices (residues 470-480 and 1045-1055) in the crRNA *vs.* tgRNA (red), tgRNA *vs.* atgRNA (gray) and atgRNA *vs.* crRNA (blue) bound systems. **B.** Projection of the JSD BB torsions on the three-dimensional structure of the HEPN1-2 catalytic core, color-coded according to the scale on the right. **C.** Distribution of the Solvent Accessible Surface Area (SASA) in the crRNA-(green), tgRNA-(violet), and atgRNA-(magenta) bound Cas13a. Vertical lines indicate the experimental values in the crRNA-(PDB: 5XWY) and tgRNA-(PDB: 5XWP) bound proteins. The statistical significance among the distributions was calculated using a two-tailed unpaired t-test (P-value reference: not significant, ns P > 0.05, * P ≤ 0.05, ** P ≤ 0.01, *** P ≤ 0.001). **D.** Representative snapshots of Cas13a bound to crRNA (top) and tgRNA (bottom), displaying a closed (crRNA-bound) and open (tgRNA-bound) conformation of the HEPN1-2 catalytic cleft.

These observations indicate that when Cas13a is bound to the crRNA alone, the HEPN1-2 catalytic cleft assumes a closed conformation. On the other hand, tgRNA binding results in an opening of the cleft. Moreover, the plasticity of the HEPN1-2 cleft appears to be a key distinguishing factor between the tgRNA- and atgRNA-Cas13a systems. In the atgRNA-Cas13a complex, the catalytic cleft is more flexible in comparison to the tgRNA-bound Cas13a, suggesting that also the length of base-pair complementarity impacts the HEPN dynamics.

### Target RNA shifts the dynamics of Cas13a

To understand how the tgRNA and an extended tag–anti-tag pair, how they bind at the level of REC lobe, and how they impact the dynamics of the spatially distant HEPN1-2 domains, it is imperative to measure the dynamic correlations among spatially distant sites (41). We employed a generalized correlation (GC) analysis, relying on Shannon’s entropy-based estimation of mutual information (35), to describe the linear and non-linear couplings of amino acids and nucleobases.

Analysis of the per-domain GC reveals that tgRNA binding increases the overall inter-domain correlations in Cas13a, including the catalytic HEPN1-2 domains, with respect to the crRNA- and atgRNA-bound complexes (**Figure 3A**). This observation reflects the biophysical basis of protein allostery (42, 43). Accordingly, the binding of an allosteric effector – in the present case the tgRNA that activates Cas13a – shifts the protein conformational landscape (30). This results in an overall increase of coupled motions, which mediate the communication between distant sites. Correlations between the crRNA and the Cas13a domains are found to improve in the ternary complexes, compared to the crRNA-Cas13a. Dynamical differences are further captured by Principal Component Analysis (PCA), which separates the binary crRNA-Cas13a complex from the ternary complexes along the first principal component (PC1), while distinguishing the tgRNA- and atgRNA-bound complexes along PC2. This observation is in line with the domain-wise PCs, which indicate that the main differentiation between the tgRNA- and atgRNA-bound Cas13a systems lies in the PC2 of the NUC domains, particularly the Linker and the HEPN1-2 domains.

**Figure 3.**
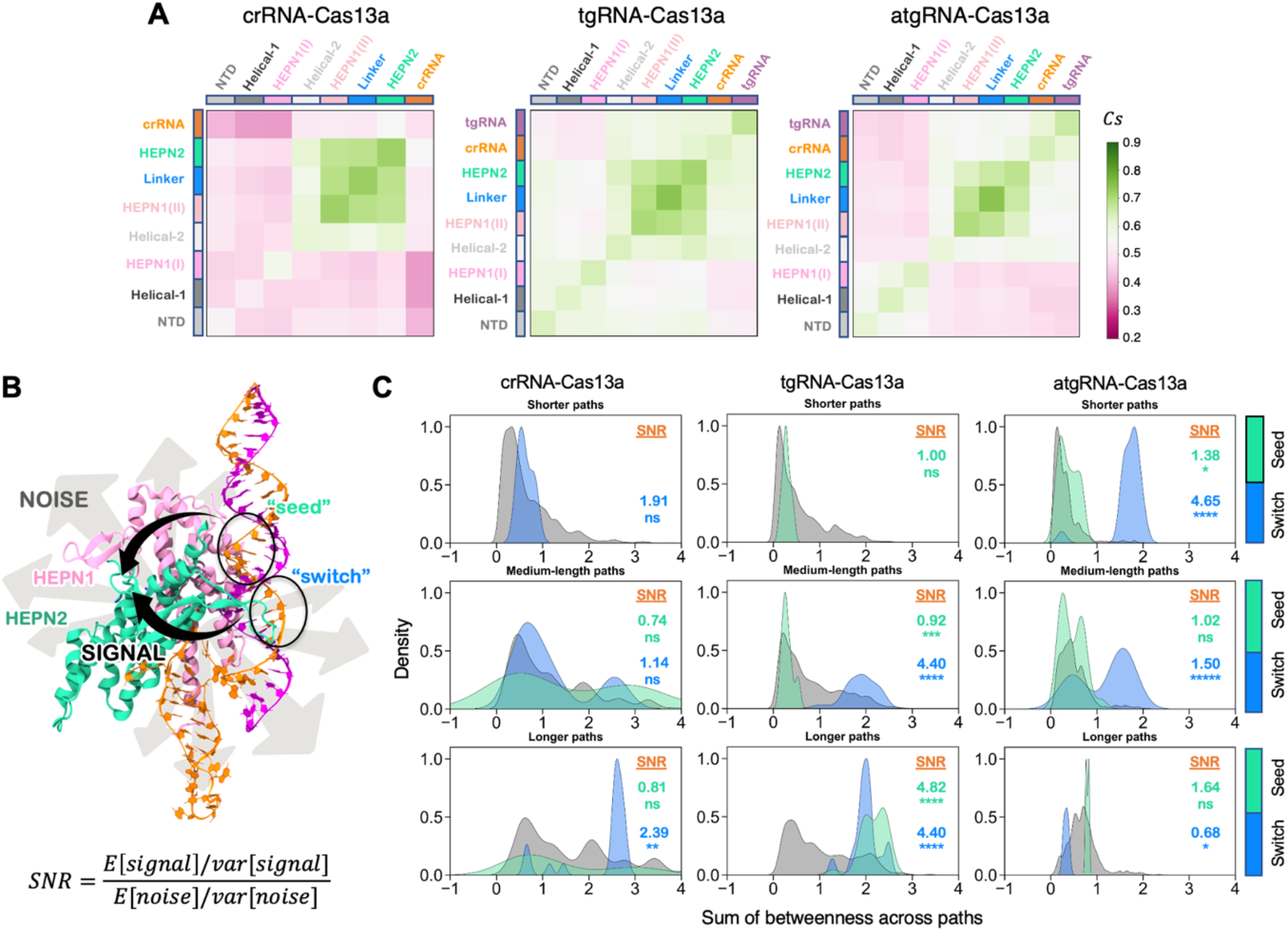
Coupled dynamics and signal-to-noise ratio of communication efficiency. **A.** Generalized inter-domain correlation maps for the crRNA-, tgRNA-, and atgRNA-bound Cas13a complexes. A pink–to–green colour code is used to describe low–to–high correlations. **B.** Schematic representation of the allosteric signals from the crRNA “seed” (nt. 9-14) and “switch” (nt. 5-8) regions to the HEPN1-2 catalytic core residues (R472, H1053, H477, R1048) over the noise, computed as the communication between all pairs of crRNA bases and Cas13a residues. Two black arrows are used to indicate the signal standing up over the noise, depicted using grey arrows. The Signal-to-Noise Ratio (*SNR*) is computed as the mean-variance ratio (*E*[]/*var*[]) of the signal over the noise (details in the *Material and Methods*). **C.** Distribution of the signals from the crRNA “seed” (green) and “switch” (blue) regions to the catalytic core residues, plotted on the background of noise (grey), across short (6-8 edge counts), medium (9–11), and long paths (12–14). The statistical significance of the signals over the noise was computed using *Z*-score statistics with a two-tailed hypothesis (P-value reference: not significant, ns P > 0.01, * P ≤ 0.01, ** P ≤ 0.001, *** P ≤ 0.0001, **** P ≤ 0.00001). Values of *SNR* are also reported for each sourcing region.

### Signal-to-Noise Ratio of communication efficiency

To understand how the observed dynamical differences impact allosteric signalling, we performed graph theory-derived network analysis (36). This approach is suited for the characterization of allosteric mechanisms, as shown in a number of studies (36, 44–46), including those performed by our research group (27–31).

We intended to estimate the communication efficiency between the crRNA spacer region and the catalytic cleft in the systems. In this respect, traditional shortest-path measurements are useful to find the most likely communication pathways between sites, but they do not report how the identified pathways stand out (i.e., are favourable) over the entire communication network (36). Assessing this favourability is crucial since the dominant allosteric pathways are particularly effective in transmitting communication between distantly coupled subunits. Hence, we introduced a Signal-to-Noise Ratio (*SNR*) estimate of the information transfer, which measures the preference of communication between predefined distant sites – i.e., the signal – over the remaining pathways of comparable length in the network – i.e., the noise. High *SNR* values indicate the preference of the network to communicate through the signal over other noisy routes (see *Material and Methods*).

First, we computed the optimal (i.e., the shortest) and top five sub-optimal pathways between all crRNA bases and the Cas13a residues, obtaining a distribution of communication efficiency in terms of the sum of edges betweennesses (i.e., the number of shortest pathways that cross the edge, measuring the “traffic” passing through them). This provided a comprehensive overview of all communications, constituting the crosstalk noise between crRNA and protein. Then, we computed the optimal and sub-optimal pathways communicating two regions of the crRNA spacer (i.e., “seed” and “switch”), which have been reported to be critical for Cas13a activation) (18), with the HEPN1-2 catalytic residues (R472, H477, R1048, H1053). This represents the signal of our interest. To ensure consistency in the scale of comparison, we characterized the *SNR* across short (6-8 edge counts), medium (9–11), and long paths (12–14), based on the number of edges communicating the crRNA regions with the catalytic residues. The obtained *SNR* indicates the extent to which the signal deviates from the distribution of noise, thus reflecting the prevalence of the signal over the noise (**Figure 3B**). Little–to–no overlap of the signal with the noise (i.e., high *SNR*) indicates the prevalence of the allosteric signal.

In the crRNA-Cas13a, broad noise distributions overlap with the signals irrespective of the path lengths, dampening the *SNR* compared to the tgRNA-Cas13a. In the tgRNA-Cas13a, amplification of signals from the “switch” regions for medium–to–long path lengths indicate that tgRNA binding improves the crosstalk efficiency between the crRNA spacer and the HEPN1-2 catalytic residues. The “seed” region also amplifies the signal across longer paths. Signals sourcing from other regions of the crRNA spacer were also computed in the tgRNA-bound complex, reporting a lower *SNR* with respect to the “switch” and “seed” regions. This observation agrees with previous biochemical data (18), suggesting that the complementarity at the “switch” region could trigger the allosteric activation of HEPN1-2, with mismatches in this region preventing LbuCas13a activation. In the atgRNA-Cas13a, a modest *SNR* is detected over medium and long path lengths, while the communication increases over shorter paths for signals sourcing from the “switch” region.

Overall, the communication is remarkably efficient in the tgRNA-bound Cas13a, in line with the experimental suggestion that tgRNA binding allosterically triggers activation of the HEPN1-2 domains (18). As noted above, allosteric phenomena are commonly associated with a shift in dynamics, resulting in the efficient transfer of signals. Our findings thereby suggest that efficient signalling is associated with a shift in correlations that prioritize the communication signals over communication noise. Therefore, shifted correlations result in noise-free communication pathways for allosteric signals.

### Key residues for allosteric coupling

The *SNR* revealed that the communication signal from the crRNA spacer to the HEPN1-2 catalytic cleft is remarkably efficient in the tgRNA-Cas13a. To further understand the signal transduction mechanism in this system, we computed the signalling pathways (i.e., the optimal and top five suboptimal paths) connecting the “switch” and “seed” regions of the crRNA to the catalytic core residues (**Figure 4A**).

**Figure 4.**
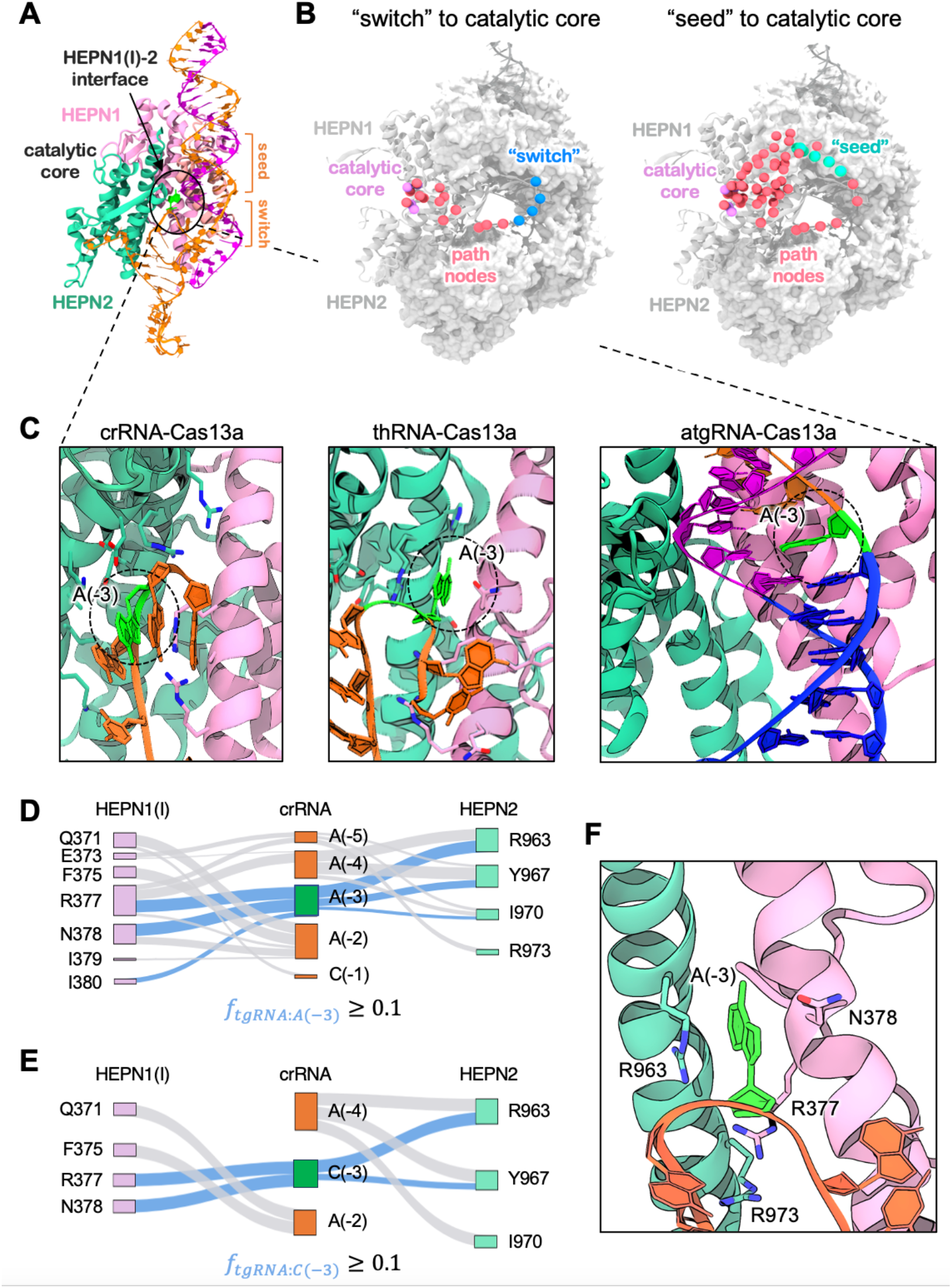
Allosteric signalling from the crRNA “switch” and “seed” regions in the tgRNA-Cas13a. **A.** Overview of the crRNA “seed” and “switch” regions binding the HEPN1(I)-2 interface, and location with respect to the catalytic core. **B.** Signalling pathways connecting the crRNA “switch” (left) and “seed” (right) regions to the HEPN1-2 catalytic residues (R472, H1053, H477, R1048). Residues occurring in the top five optimal pathways of communication are plotted on the three-dimensional structure of CRISPR-Cas13a (grey). Residues are plotted using spheres of different colours according to the region of interest: “switch” (blue), “seed” (green), catalytic residues (pink), and remaining path residues (red). **C.** Conformation of the A(−3) base (green) of the crRNA repeat region, in proximity to the HEPN1(I)-2 interface of the three RNA-bound complexes. **D-E.** Sankey plots reporting the frequency, *f*, of formed contacts between residues of the HEPN1(I) and HEPN2 domains, and the crRNA, forming for ≥ 10 *i* of the simulation time (*f* ≥ 0.1) in the tgRNA-Cas13a (D), and in the system substituting the A(−3) base with cytosine (E). Residue pairs of HEPN1(I) (left), HEPN2 (right), and the crRNA bases (centre) are connected by edges whose width is proportional to *f*. Contact edges that involve A/C(−3) are shown in blue, to highlight them with respect to the remaining interactions (grey). **F.** Close-up view of the HEPN1(I)-2 interface in the tgRNA-bound Cas13a. Four polar/positively charged residues (R377, N378, R963, and R973; cyan) have been mutated in alanine to explore their impact on Cas13a activity.

The pathways connecting the “switch” region to the catalytic residues exclusively follow a route that directly connects the crRNA bases of the repeat region (A(−5)–C(−1)) to the catalytic core through the HEPN1(I)-2 interface (**Figure 4B**). On the other hand, the “seed” region communicates with the catalytic core through multiple routes, involving the Linker-HEPN2 interface, the HEPN1(II) residues, as well as the HEPN1(I)-2 interfacial residues through the crRNA repeat, similar to the communication observed for the “switch”.

Hence, the pathways connecting the “switch” nucleotides to the catalytic residues display a lower number of residues with increased occurrences, tracing a more efficient communication path, compared to the pathways connecting the “seed”. This observation further affirms the critical role of the “switch” in the allosteric activation of HEPN1-2 (18). Extending our analysis to 50 suboptimal pathways reported a similar trend, where a direct route consistently connects the “switch” region to the HEPN1(I)-2 catalytic core through the crRNA repeat bases A(−4) and A(−3). This evidence pinpoints a pivotal role of the crRNA repeat region in signal transmission.

To better understand our observations, we analysed the interactions between the crRNA repeat region (A(−5)–(C-1)) and the proximal HEPN1(I)-2 interface (residues: 371–383; 963– 975). Notably, this interface region is located distally with respect to the catalytic cleft (**Figure 4A**). We observed that in the tgRNA-bound system, the A(−3) base of the crRNA repeat region penetrates the interface (**Figure 4C**), hampering interactions between HEPN1(I) and HEPN2. This is confirmed by the histogram of differential contact stability (Δ*f*_B;C-D13?C-D_), showing that, in the crRNA-Cas13a, the flipped-out A(−3) base causes increased interactions at the HEPN1(I)-2 interface, with respect to the tgRNA-bound system. In the atgRNA-Cas13a, the crRNA repeat region is sequestered due to the extended tag–anti-tag complementarity, leading to stable interfacial contacts compared to the tgRNA-Cas13a.

To detail the interactions of the A(−3) base in the tgRNA-bound Cas13a, we conducted an in-depth contact analysis at the HEPN1(I)-2 interface. A Sankey plot was used to report the frequency, *f*, of stable contacts between residues of the HEPN1(I) and HEPN2 domains, and the crRNA, forming for ≥ 10 *i* of the simulation time (*f* ≥ 0.1) in the tgRNA-Cas13a (**Figure 4D**).

In this plot, residues are connected through edges, whose width is proportional to *f*. We observe that A(−3) substantially interacts with polar/positively charged residues, mainly R377, N378, and R963. Analysis of the differential contact stability was also carried out to characterize interactions that gain stability in tgRNA-Cas13a, compared to the crRNA-Cas13a.

As expected, a loss of stable contacts at the HEPN1(I)-2 interface is observed in the tgRNA-Cas13a, compared to the crRNA-bound crRNA system. Upon tgRNA binding, R377 and N378 of HEPN1(I) gain interactions with A(−3); and R963 and R973 of HEPN2 increase their contacts with the neighbouring bases, compared to the crRNA-Cas13a. To test whether this observation is sequence dependent, we substituted the A(−3) base with a smaller cytosine in the tgRNA-Cas13a and carried out an additional ∼5 μs-long simulation. In this system, the C(−3) base stably locates at the HEPN1(I)-2 interface and interacts with R377, N378, and R963 (**Figure 4E**). Taken together, these observations suggest that these rearranged interactions involving charged/polar residues between HEPN1(I)-2 could be critical for allosteric signalling from the crRNA spacer to the catalytic core.

### Role of HEPN1-2 interfacial interactions

To experimentally verify our observations, we generated and purified the wild-type (WT) Cas13a and four variants, mutating charged/polar residues to alanine at the HEPN1(I)-2 interface (i.e., R377A, N378A, R963A, and R973A, **Figure 4F**). We designed and generated tgRNAs and crRNAs containing the spacer used by Liu et al.(19) (PDB: 5XWP, **Figure 5A**) and an anti-tag containing target RNA (atgRNA), holding an extended 8-nt. sequence with complementarity to the crRNA direct repeat (**Figure 5B**). We performed fluorescent RNA trans-cleavage assays with WT LbuCa13a and the variants bound to our tgRNA and atgRNA sequences (see *Material and Methods*).

**Figure 5.**
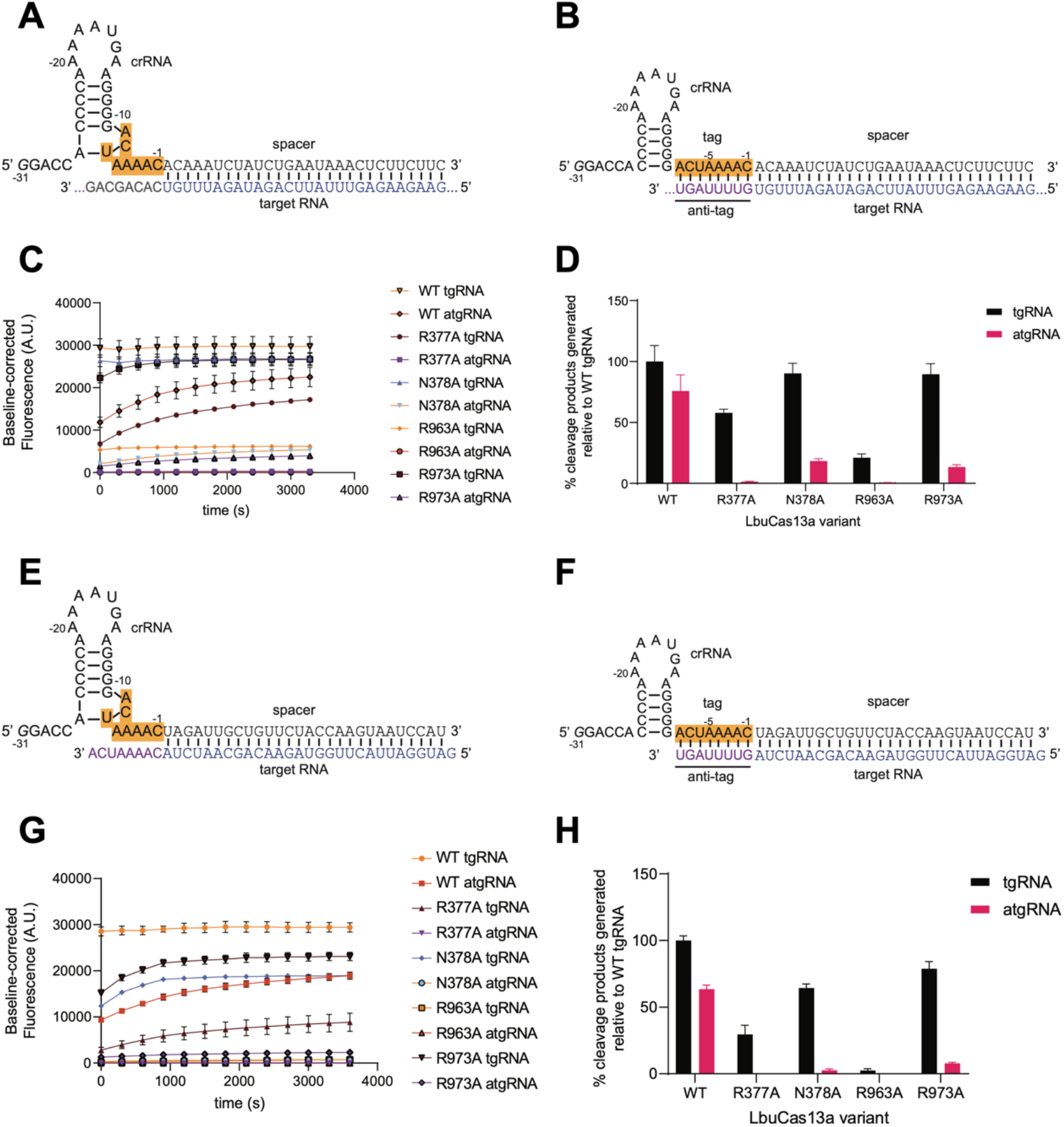
LbuCas13a variants are more sensitive to inhibition by anti-tag-containing RNAs. **A-B.** crRNA-target RNA (tgRNA) pair (A) and crRNA-anti-tag RNA (atgRNA) pair (B), designed based on Liu et al. (19). **C.** One-hour time course of background-corrected fluorescence measurements from RNA cleavage experiments by LbuCas13a WT and variants initiated by the addition of 100 pM tgRNA or atgRNA shown in (A) and (B). **D.** Data from (C) normalized as percent cleavage product generated relative to WT LbuCas13a with tgRNA. **E-F.** crRNA-target RNA (tgRNA) pair (E) and crRNA-anti-tag RNA (atgRNA) pair (F), designed based on Wang et al.(20) **G.** One-hour time course of background-corrected fluorescence measurements from RNA cleavage experiments by LbuCas13a WT and variants initiated by the addition of 100 pM tgRNA or atgRNA shown in (I) and (F). **H.** Data from (G) normalized as percent cleavage product generated relative to WT LbuCas13a with tgRNA.

In the WT LbuCas13a, there is robust activation of the nuclease activity with a tgRNA, while the presence of an atgRNA results in a small decrease in the apparent cleavage rates over time (**Figure 5C**), albeit the amount of end-point cleavage product was only slightly reduced relative to the tgRNA (**Figure 5D**). In our LbuCas13a variants, N378A and R973A maintained a robust activity for the tgRNA, the R377A variant suffered a reduction in cleavage efficiency and R963A was not active at all (**Figure 5D**). In the presence of an atgRNA, none of the variants displayed any significant nuclease activation, suggesting that these variants are more sensitive to anti-tag containing RNAs than the WT LbuCas13a (**Figure 5C-D**). To further validate these observations, we obtained crRNA-tgRNA (**Figure 5E**) and crRNA-atgRNA pairs (**Figure 5F**) corresponding to the sequences used by Wang et al. (PDB:7DMQ) (20). Our cleavage assays showed a similar activation pattern to our data above, where our LbuCas13a variants show increased inhibition in the presence of an atgRNA (**Figure 5G-H**).

Taken together, these experimental data, using a 28 nt. spacer similar to the simulations, reveal that the WT LbuCas13a maintains its activity irrespective of tgRNA or atgRNA binding. On the other hand, our variants still cleave RNA with tgRNA bound, while hampered nuclease activation in the presence of an extended tag–anti-tag complementarity is observed.

### Cas13a variants reorganize the allosteric communication network

To evaluate how the signalling transfer is affected by the mutations we made, we computed the *SNR* on a ∼15 μs ensemble for each of our four Cas13a variants bound to a tgRNA and atgRNA and compared it to the WT Cas13a. We compared the maximal *SNR* detected in the variants with that of the WT Cas13a, considering all signals sourcing from the critical “switch” region, and establishing a consistent scale for comparison (**Figure 6A**). The highest *SNR* across different path lengths indicates the most favoured communication route in the system, irrespective of path lengths. We thereby computed the ratio between the maximal *SNR* in the variants and in the WT Cas13a (*SNR*_;:3*i*E_= *SNR*_F:G1H:;*i*:03_ /*SNR*_F:G1I3_). This comparison indicates whether point mutations at the HEPN1(I)-2 interface impact the strength of the communication between the crRNA “switch” and the catalytic residues with respect to the WT.

**Figure 6:**
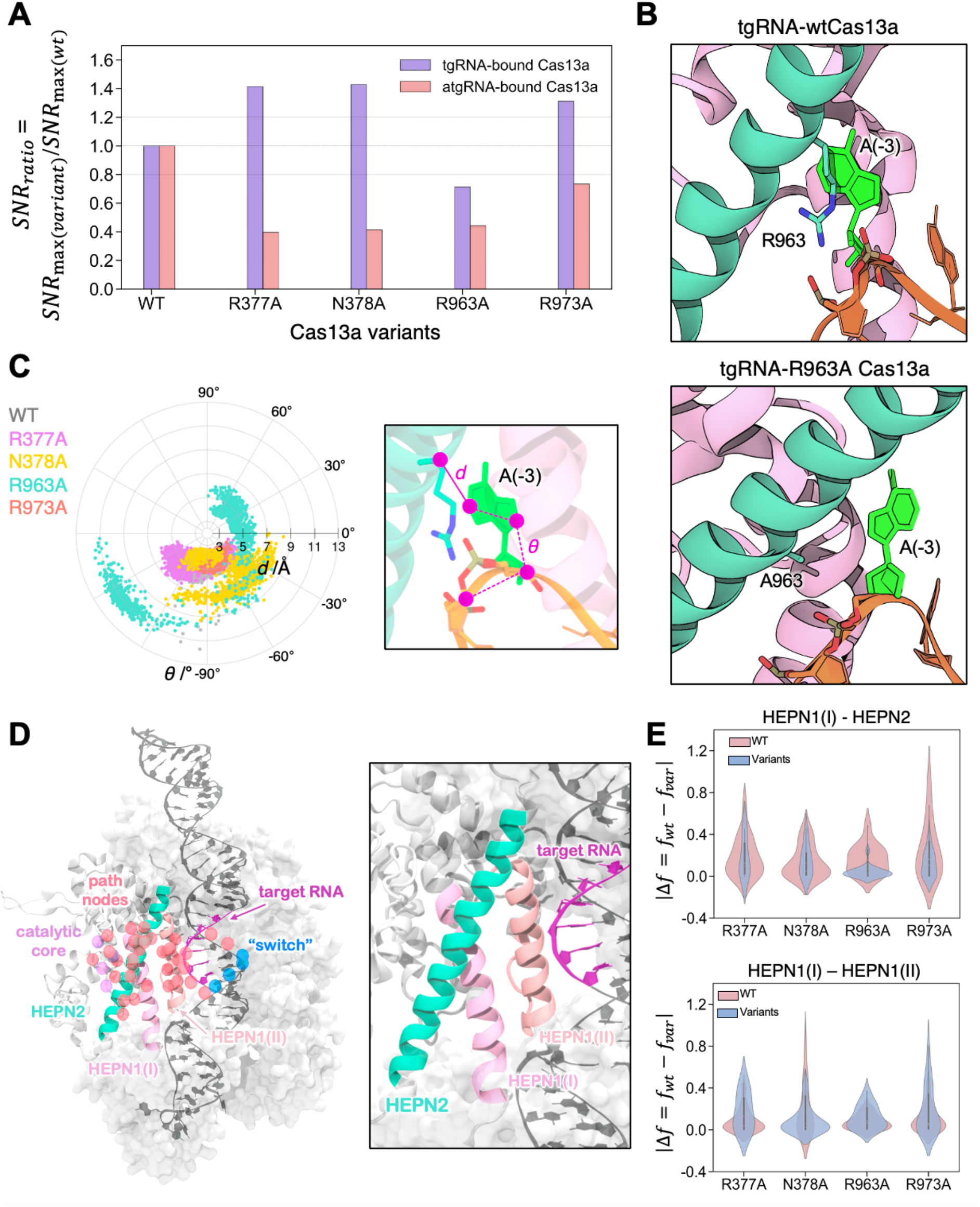
Mutation-induced reorganized communication network. **A.** Ratio between the maximal Signal-to-Noise Ratio (*SNR*_F:G_) in our Cas13a variants and the WT Cas13a (*SNR*_;:3*i*E_ = *SNR*_F:G1H:;*i*:03_/*SNR*_F:G1I3_), computed for the tgRNA-(violet) and atgRNA-(mauve) bound systems. **B.** Close-up view of the HEPN1(I)-HEPN2 interface in the WT tgRNA-Cas13a (top) and its R963A mutant (bottom), showing the extrusion of A(−3) from the protein in the R963A mutant. **C.** Polar plot of (*i*) the distance *d* between the C⍺ atom at position 963 and the centre of mass of the N1-C6 ring (polar coordinate, in Å), and (*ii*) the dihedral angle *θ* between the C3’@A(−4), C3’@A(−3), C8@A(−4) and C2@A(−4) atoms (angular coordinate, in degrees), computed from the simulated ensembles of the WT tgRNA-bound Cas13a and its mutants. The *d* distance and *θ* angle are shown on the right. **D.** Signalling pathways connecting the crRNA “switch” region to the HEPN1-2 catalytic residues (R472, H1053, H477, R1048) in the atgRNA-Cas13a. A close-up view of the HEPN1(I), HEPN1(II) and HEPN2 interfaces involved in the allosteric pathways is reported on the right. **E.** Distributions of differential contact stability at the HEPN1(I)-HEPN2 (top) and HEPN1(I)-HEPN1(II) (bottom) interfaces, between the atgRNA-Cas13a and its variants (|Δ*f* = *f*_I3_ − *f*_H:;_|).

We observe that in the tgRNA-bound systems, the R377A, N378A, and R973A variants maintain a high *SNR*_;:3*i*E_, as evidence of efficient communication compared to the WT Cas13a (**Figure 6A**). On the other hand, R963A reduces the signal with respect to the WT, indicating altered communication.

Upon atgRNA binding, the *SNR*_;:3*i*E_reaches 40-60 *i* of perturbations in all variants with respect to the WT, with R973A maintaining a *SNR*_;:3*i*E_ approximately within 30 *i* of the WT. This suggests that, in the atgRNA-bound variants, the signalling from the “switch” to the HEPN1-2 catalytic core is reduced. This is in line with the experimental activity, showing that none of the variants displays a significant nuclease activation in the presence of an atgRNA (**Figure 5C-D**).

To further understand the signalling transfer in the tgRNA-bound variants, and the observation of a reduced *SNR*_;:3*i*E_ in the inactive R963A mutant, we analysed the interactions of the crRNA repeat bases and the proximal HEPN1(I)-2 interface. In the WT tgRNA-Cas13a, the A(−3) base forms stable contacts with R377, N378, and R963, along with several other HEPN2 residues (**Figure 4D**). These interactions are preserved in the tgRNA-bound mutants except, as expected, for the mutated residue. Nevertheless, in the tgRNA-bound R963A mutant, the A(−3) base is more flexible and is frequently extruded from the HEPN1(I)-2 interface (**Figure 6B**). The A(−3) conformations are monitored on a polar plot reporting the distance *d*, which describes the displacement of A(−3) with respect to the C⍺ atom at position 963, and the dihedral angle *θ*, reporting the rotation of the A(−3) purine base with respect to the crRNA backbone (**Figure 6C**). The polar plot evidences the higher flexibility of A(−3) in the R963A mutant, compared to the remaining systems, and its extrusion from the HEPN1(I)-2 interface. This observation can be ascribed to the loss of interaction between the R963 guanidinium side chain and the crRNA phosphate backbone that, on the other hand, is maintained in the other systems. We recall that the R963A substitution hampers the tgRNA-Cas13a activity (**Figure 5**). This suggests that the positioning of the A(−3) base, and the dynamics of the crRNA repeat region at the HEPN1(I)-2 interface, critically affects the transmission of the signal from the “switch” to the catalytic core, as evidenced by lower *SNR* with respect to the WT.

To characterize the allosteric communication in our atgRNA-bound variants, and how it compares to the tgRNA-Cas13a, we inspected the pathways communicating the “switch” to the HEPN1-2 catalytic core. In the atgRNA-bound WT Cas13a, allosteric pathways mainly involve the HEPN1(I), HEPN1(II), and HEPN2 interfaces and the tgRNA (**Figure 6D**), at odds with the direct routes passing through the crRNA repeat and HEPN1(I)-2 observed in the tgRNA-Cas13a (**Figure 4**). When introducing our point mutations in the tgRNA-bound Cas13a, the major routes of communication are similar to those of the WT system, with the addition of a few residues from HEPN1(II). In the atgRNA-bound variants, allosteric pathways are more sensitive to perturbations, compared to the tgRNA-bound variants. In the presence of an atgRNA, the variants increase the number of crRNA bases involved in the communication with respect to WT atgRNA-Cas13a, with the N378A and R963A variants also losing the tgRNA communication channel. An analysis of the interactions at the major interfaces along the allosteric pathways (i.e., HEPN1(I), HEPN1(II) and HEPN2, **Figure 6E**) also shows that the mutations result in a reduction of stable contacts at the HEPN1(I)-HEPN2 interface with respect to the WT Cas13a, while gained back at the HEPN1(I)-HEPN1(II) interface. These rearrangements at the critical HEPN1(I)-(II)-HEPN2 interfaces are the basis of the altered signalling pathways observed in the atgRNA-bound variants.

In summary, analysis of the allosteric communication in our mutants reveals that the inactive mutants reduce the signal with respect to the WT. This altered communication can be attributed to the dynamics of the crRNA repeat region in the tgRNA-bound systems, and to an overall perturbation of the crosstalk pathways in the presence of an atgRNA.

## DISCUSSION

Here, extensive MD simulations were combined with graph theory and experimental assays to characterize the allosteric activation mechanism in Cas13a, a newly emerged CRISPR-Cas system, which is being developed as a powerful tool for RNA cleavage, detection and imaging.

Multiple-μs simulations reveal that the binding of a target RNA (tgRNA) at the recognition lobe impacts the dynamics of the spatially distant HEPN1-2 catalytic core, resulting in altered dynamics with respect to the crRNA-bound form (**Figure 2A-B**), and in the opening of the catalytic cleft (**Figure 2C**). This is in line with biochemical data, suggesting that tgRNA binding allosterically activates HEPN1-2 to form a composite active site (15, 18). In the presence of an extended tag–anti-tag pairing (atgRNA), the HEPN1-2 catalytic cleft increases its flexibility, compared to both the crRNA- and tgRNA-Cas13a, indicating that the length of the complementarity also affects the HEPN1-2 dynamics.

Analysis of mutually coupled motions shows that tgRNA binding induces a shift in the system’s dynamics (**Figure 3A**), evidenced by increased inter-domain correlations. Coupled dynamics are also found in the presence of an atgRNA, mainly involving the crRNA and the Cas13a domains. This “shift in dynamics” is typical evidence of an allosteric response (30, 42), reinforcing the notion of the tgRNA as an effector of Cas13a’s allosteric function.

To better understand how the observed differences in the systems’ dynamics impact the flux of information from the sites of tgRNA binding to the catalytic core, we estimated the crosstalk efficiency across the studied systems. We analysed how the communication signal from the crRNA (site of tgRNA binding) to the HEPN1-2 cleft emerges over the remaining pathways (i.e., the noise) through a signal-to-noise ratio (*SNR*) measure. We found that, upon tgRNA binding, the signal from the “switch” region of the crRNA amplifies over the noise, as evidence of efficient crosstalk (**Figure 3B**). The communication sourcing from the “switch” is also maintained upon atgRNA binding. These observations underscore the critical role of this region in the allosteric signalling between the sites of effector binding and RNA cleavage. We recall that, in line with our observations, biochemical data have noted the “switch” region to be crucial in triggering the allosteric activation of HEPN1-2 (18).

Analysis of the allosteric pathways also shows that the “switch” region efficiently communicates with the HEPN1-2 catalytic core through the crRNA bases of the repeat region (i.e., A(−5)-C(−1), **Figure 4B**). This pinpoints a cardinal role for the crRNA repeat region, which prompted us to analyse its interactions with Cas13a. We looked at the interactions with the proximal HEPN1(I)-2 interface that, notably, locates distally with respect to the catalytic cleft (**Figure 4C**). We observe that in the tgRNA-Cas13a, the A(−3) base of the crRNA repeat penetrates the HEPN1(I)-2 interface, decreasing inter-domain interactions compared to the crRNA-Cas13a where A(−3) is extruded. The extended tag–anti-tag complementarity also sequesters A(−3) from the HEPN1(I)-2 interface. The interaction network further reveals several polar/charged residues critical for HEPN(I)-2 binding, that exhibit increased interactions with A(−3) upon tgRNA biding (**Figure 4D-E**). In detail, N378, R963A, R973, and R377A rearrange their interactions at the HEPN(I)-2 interface upon tgRNA binding, which could be critical in mediating the allosteric information transfer process.

To experimentally test the role of these polar/charged residues in the allosteric activation of Cas13a, we performed mutagenesis and RNA cleavage experiments. Alanine substitution of N378, R377, and R973 maintains a robust activity for the tgRNA, while R963A is not active (**Figure 5**). In the presence of an atgRNA, none of the variants displays significant nuclease activation. This reveals that our N378A, R377A, and R973A variants can discriminate tgRNA for cleavage over atgRNA. Our computational analysis and the experimental assays thereby disclose polar/charged residues that are pivotal for the allosteric signalling and whose mutation in alanine can push the preference for tgRNA-activated cleavage vs. atgRNA-activated cleavage.

To understand the mechanistic function of our mutants, we performed additional multi-μs long simulations of our variants. Analysis of the allosteric signal sourcing from the “switch” region to the HEPN1-2 catalytic cleft, shows that our variants mostly perturb the *SNR* upon atgRNA binding (with respect to the WT atgRNA-Cas13a, **Figure 6A**), while the tgRNA-bound variants preserve efficient signalling compared to the WT tgRNA-Cas13a. Interestingly, the tgRNA-bound R963A Cas13a, which is experimentally shown to be inactive (**Figure 5D**), reduces the signal with respect to the WT, indicating altered communication. In this system, the A(−3) base is frequently extruded from the HEPN1(I)-2 interface (**Figure 6C**), suggesting that the positioning of the A(−3) base (and the dynamics of the crRNA repeat region) critically affects the transmission of the signal from the “switch” to the catalytic core, as evidenced by lower *SNR* with respect to the WT.

Analysis of the allosteric pathways also shows that our variants preserve the communication connecting the “switch” to the catalytic cleft in the tgRNA-bound system. On the other hand, in the presence of a tag–anti-tag, our mutations perturb and reduce the conservation of the signalling pathways observed in the WT atgRNA-Cas13a. This observed perturbation of the allosteric crosstalk pathways, observed in the atgRNA-bound variants, along with altered *SNR* compared to the WT, is qualitatively consistent with the experimentally observed inactivity of the atgRNA-bound variants (**Figure 5**). Overall, the inactive mutants (i.e., the atgRNA-bound variants and the tgRNA-bound R963A) reduce the signal with respect to the WT. This reduced signalling can be ascribed to the dynamics of the crRNA repeat region in the tgRNA-bound systems, and to an overall perturbation of the crosstalk pathways in the presence of an atgRNA.

## CONCLUSIONS

In summary, our study characterizes the allosteric activation mechanism of Cas13a and discloses critical point mutations able to promote tgRNA-mediated over an extended tag-anti-tag-mediated complementarity, for RNA cleavage activation. We show that the binding of a tgRNA acts as an allosteric effector of the spatially distant HEPN1-2 catalytic cleft, by amplifying the allosteric signals over the dynamical noise. The allosteric signal connecting the sites of tgRNA binding with the HEPN1-2 catalytic site stands up over the dynamical noise sourcing from the crRNA “switch” region, and communicating through the crRNA repeat, whose A(−3) base regulates interactions at the buried HEPN1(I)-2 interface. Critical residues in this region, R377, N378, and R973 rearrange their interactions upon tgRNA binding and are shown experimentally to select tgRNA, over an extended atgRNA, for RNA cleavages. Computational analysis of our Cas13a variants also reveals that the inactive mutants reduce the signal compared to the WT system, offering a rationale that is in line with the allosteric functioning of Cas13a, proposed experimentally (15, 18). Finally, considering the selectivity of the mutants toward tgRNA over an extended complementary beyond the guide-target duplex, we speculate here that alanine mutation of R377, N378, and R973 could improve (or alter) the selectivity of Cas13a. This hypothesis is confirmed in our companion paper by Molina-Vargas et al. (*under review*). Taken together, our findings offer a fundamental understanding of the CRISPR-Cas13a mechanistic function, and pave the way for the development of innovative RNA-based cleavage and detection tools.

## AUTHOR CONTRIBUTIONS

**Souvik Sinha**: Conceptualization, Data curation, Formal Analysis, Investigation, Methodology, Validation, Visualization, Writing-original draft. **Adrian M. Molina Vargas**: Data curation, Investigation, Formal Analysis, Writing—review *j* editing. **Pablo R. Arantes**: Formal Analysis, Methodology, Validation, Visualization, Writing—review *j* editing. **Amun Patel**: Formal Analysis, Methodology, Validation, Visualization, Writing—review *j* editing. **Mitchell R. O’Connell**: Supervision, Project administration, Funding acquisition, Writing—review *j* editing. **Giulia Palermo**: Conceptualization, Supervision, Project administration, Funding acquisition, Writing-original draft.

